# Exploring the intrinsic and extrinsic determinants of heterogeneity in a *β*-cell network

**DOI:** 10.1101/2025.10.26.684303

**Authors:** Anwar Khaddaj, Shane Browne, Woori Choi, Vira Kravets, Andrew G. Edwards

## Abstract

Pancreatic islets are micro-organs composed of multiple endocrine cell types. *β*-cells are the most common of these and are highly heterogeneous in both their intrinsic properties, such as ion channel conductances and metabolic activity, and extrinsic properties, including gap junction coupling and paracrine signaling. Capturing these diverse sources of heterogeneity is essential for computational models that aim to reproduce islet function. We evaluate two established multicellular models, the coupled Cha-Noma model of the mouse *β*-cell network, and the Riz human model. Prior work with these two models suggest that the Cha-Noma model involves minimal intrinsic heterogeneity and therefore bursts with highly synchronized activity, whereas heterogeneity in the Riz model prescribed from *in vitro* patch-clamp results in highly unsynchronized behavior of the coupled network. We hypothesize that adjusting the number of bursting cells in both model formulations may invoke more physiologically realistic network coordination. We applied a categorical sensitivity analysis to establish which parameters are most important for determining bursting in single cells of the Riz model. We also introduced new heterogeneity (based on single-cell gene expression) in parameters that had previously been treated as invariant among *β*-cells. We hypothesized that introducing heterogeneity in the small conductance Ca^2+^ K^+^ channel in particular could promote a higher proportion of bursting cells. Lastly, both models fail to incorporate the influences of non-*β*-cells. We introduced paracrine signaling between *α* and *β*-cells into the coupled Cha-Noma model and showed it plays a role in accelerating the response time of *β*-cells to acute glucose stimulation (i.e. promotes a 1^st^ responder phenotype).

## 5.1 Introduction

The Islet of Langerhans is a ‘micro-organ’ consisting of several hormone-secreting cells. These endocrine cells include insulin-secreting *β*-cells, glucagon-secreting *α*-cells, and somatostatin-secreting *δ*-cells. The predominant cell type in both human and mouse islets are *β*-cells [1, 2, 3, 4, 5]. *β*-cells have been shown to have significant heterogeneity in both intrinsic and extrinsic parameters. This intrinsic heterogeneity includes variation in ion channel conductances [6] and in metabolic activity [7, 8, 9, 10]. Furthermore, heterogeneity in extrinsic characteristics has been broadly observed and is altered in disease, particularly type II diabetes. *In situ* patch-clamp of peripheral islet cells [11] *In vitro* patch-clamp of dissociated *β*-cell pairs [12], as well as non-invasive fluorescence recovery after photobleaching experiments [13], have shown significant heterogeneity in gap junction coupling between *β*-cells. Additionally, extrinsic heterogeneity in paracrine signaling is likely to exist within the islet due to non-uniformity in heterotypic contacts between cells that are able to engage in paracrine signaling over short distances [14, 15, 1].

For computational models of islets to be informative, there must be a proper reconstruction of both these intrinsic and extrinsic forms of heterogeneity. Published computational *β*-cell models have attempted to reconstruct this heterogeneity, and here we interrogate two of them: (1) the coupled Cha-Noma mouse model [16]; and, (2) a multicellular version of the Riz human model [17, 18]. Importantly, cursory inspection of glucose-dependent activity in both of these models indicates that neither captures function that is truly representative of intact *β*-cell networks (Figure 5.2). The coupled Cha-Noma model exhibits activity (indicated by intracellular Ca^2+^) that is much more coordinated and synchronous than is observed in experiments, while the multicellular Riz model is excessively dyssynchronous and uncoordinated. By quantifying the glucose-induced excitability of the *β*-cell populations used (i.e. the effects of intrinsic heterogeneity) in both models (Figure 5.1), we observed that the coupled Cha-Noma model had a much higher proportion of bursting cells compared to the multicellular Riz model. Thus, we first hypothesized that redressing this difference by modifying bursting-specific intrinsic heterogeneity of the cell population in the multicellular Riz model might rectify this coordination issue.

**Fig. 5.1:**
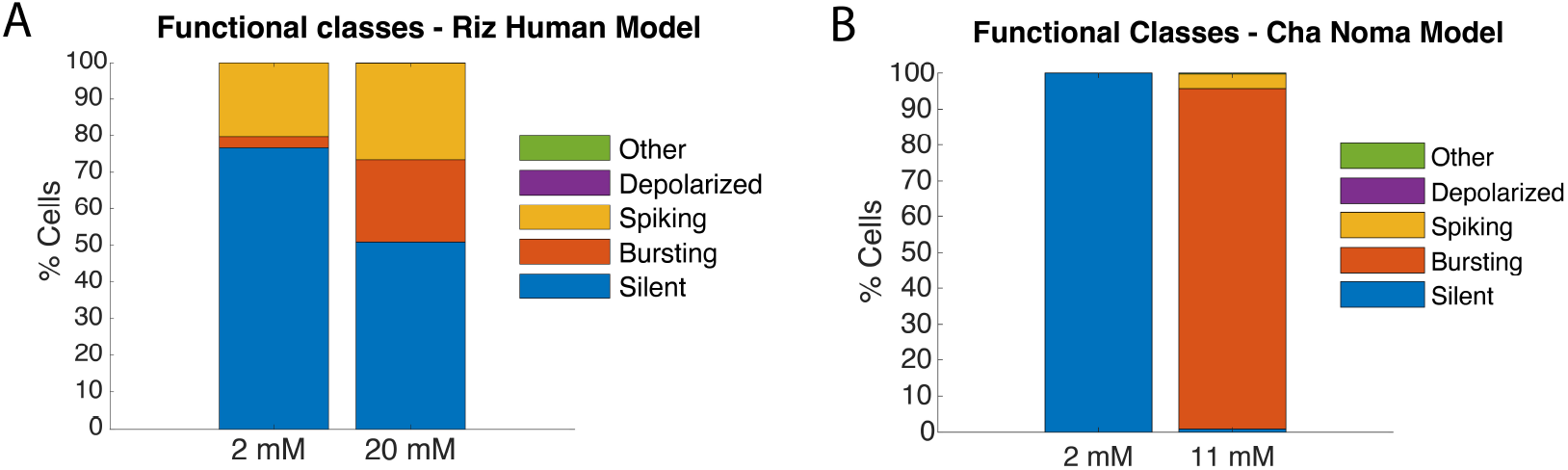
Classification of computation models into different excitability phenotypes. (A) Human Riz model with heterogeneous parameters. (B) Mouse Cha-Noma model with heterogeneous parameters.

**Fig. 5.2:**
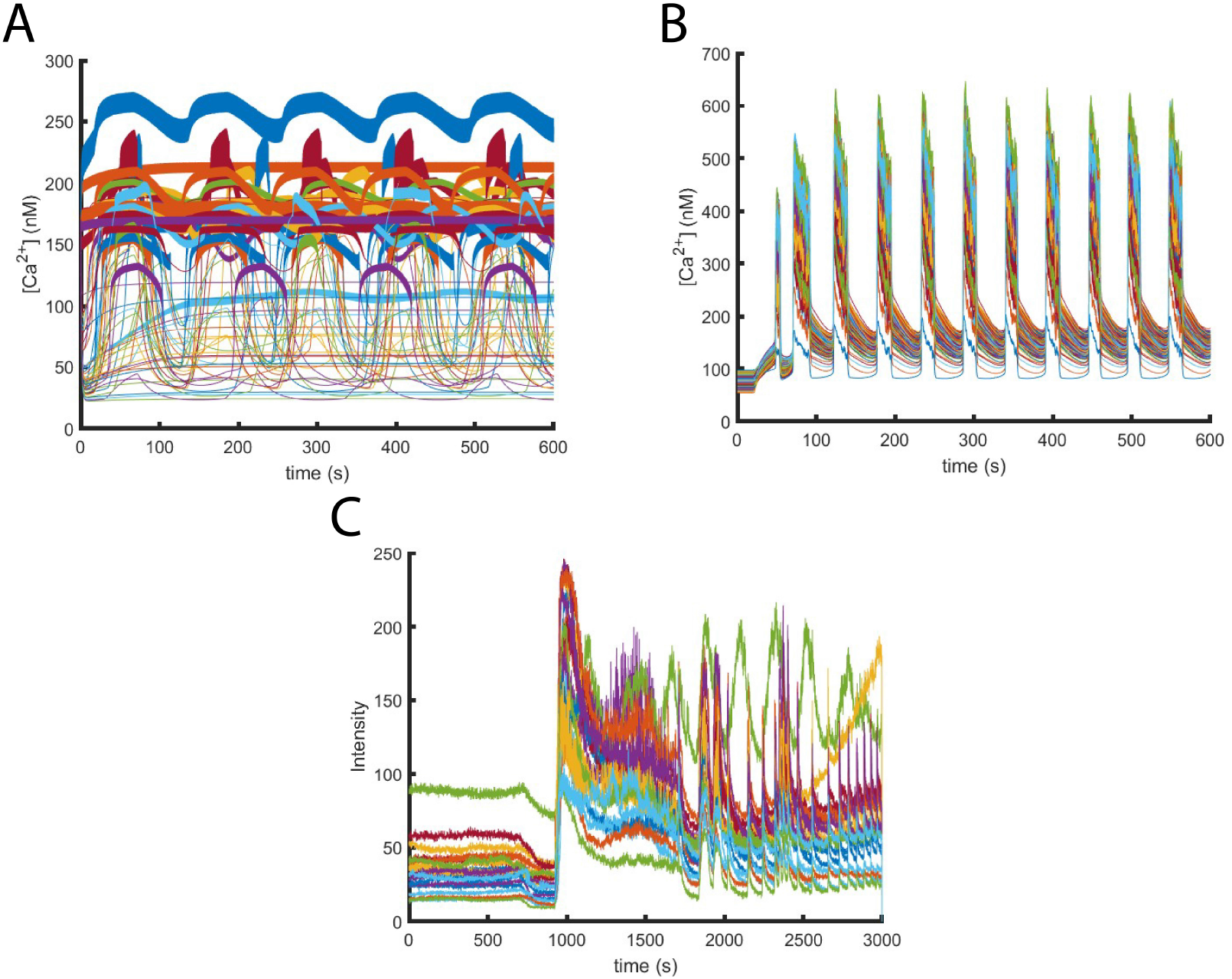
Computational and experimental calcium traces. (A) Calcium traces for 60 cells in the coupled Riz model at 20 mM glucose. (B) Calcium traces for 1000 cells in the coupled Cha-Noma model when excited from 2 mM to 11 mM glucose at t = 20 s. (C) Experimental calcium traces from an isolated mouse islet when excited from 2 mM to 11 mM glucose.

Moreover, though both computational models introduce heterogeneous parameters, not all intrinsic parameters are heterogeneous in the population due to limited availability of human-specific patch-clamp data. In the multicellular Riz model, one such parameter is the maximum conductance of the small-conductance calcium-activated potassium (SK) channel. Small-conductance calcium-activated potassium (SK) channels are key regulators of pancreatic *β*-cell excitability. Activated by elevations in intracellular calcium, SK channels generate after-hyperpolarizing currents that contribute to membrane repolarization and burst termination. By shaping oscillatory activity, SK channels influence the frequency and duty cycle of *β*-cell bursts, thereby indirectly modulating insulin secretion through their impact on calcium oscillations and granule exocytosis [19]. The assumption of uniform conductance across the population oversimplifies *β*-cell electrophysiology and fails to account for molecular variability that may be physiologically important for network function. Single-cell transcriptomic profiling of human pancreatic islets in [20] can provide an approximation of ion channel expression at the resolution of individual cells. Our reanalysis of the dataset presented by Segerstolp et al.[20], indicates that KCNN3 transcripts, encoding SK subtype 3, are consistently expressed at quantifiable levels in *β*-cells, whereas other SK isoforms (KCNN1, KCNN2, KCNN4) are barely detectable. These results suggest that KCNN3 transcript expression constitutes the main detectable SK channel transcript in human *β*-cells, and we hypothesized that introducing variability defined by cell-to-cell KNN3 transcript level may provide a new basis for modulating bursting across the modeled Riz *β*-cell population.

Lastly, it is notable that both the coupled Cha-Noma model and the coupled Riz model only include *β*-cells. Consequently, only intrinsic heterogeneity and extrinsic heterogeneity in the form of gap junction coupling are accounted for. However, paracrine signaling is another important determinant of islet function, and particularly regulation of the *β*-cell network. Application of exogenous glucagon and glucagon-like-peptide-1 (GLP-1) has been shown to potentiate glucose-stimulated insulin secretion (GSIS), and there is evidence to suggest that endogenous production of glucagon and GLP-1 from *α*-cells acts on *β*-cells via paracrine signaling through the GLP-1R [21, 22, 23]. Thus, *α*-*β*-cell communication is an important determinant of islet function. This paracrine signaling might have further implications for *β*-cell subpopulation development, as it has been hypothesized that the *β*-cell subpopulation termed 1^st^ responders is preferentially located near *α*-cells. [24, 25].

In this work, we use sensitivity analysis to help reconstruct realistic network coordination by varying the most heterogeneous parameters in the multicellular Riz model that are the most sensitively associated with the bursting phenotype. We also incorporate SK conductance heterogeneity into simulations in order to better reflect physiological diversity of SK expression in *β*-cell populations. Finally, we set out to introduce a model for *α*-*β* cell communication into the coupled Cha-Noma model, and quantify the influence of this signaling on indicators of 1^*st*^ responder behavior near *α* cells.

## 5.2 Methods

### 5.2.1 Sensitivity analysis

To study the intrinsic heterogeneity of *β*-cells within an islet, we examine how variations in model parameters influence the responses of *β* cells to elevated glucose. To achieve this, we employ a sensitivity analysis technique to quantify the impact of heterogeneous parameters on model outputs. Our study focuses on a human islet population of 3000 cells from which a subset of *n*_*b*_ = 774 *β*-cells is randomly sampled. We utilize the *β*-cell Riz model described in [26], which has *n*_*x*_ = 15 state variables and *n* _*p*_ = 86 parameters and can be formulated abstractly as an ordinary differential equation (ODE):

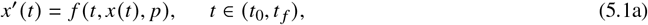

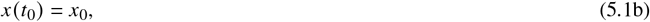

where 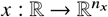 represents the state variables, 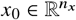 represents the initial conditions, 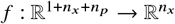, and 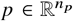 represents the parameters. Out of the 86 parameters, only 16 parameters are varied during the simulations, and these are grouped into a parameter vector 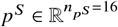:

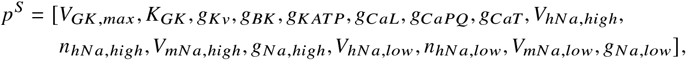

The descriptions of these heterogeneous parameters are provided in Table 5.1 below.

**Table 1:**
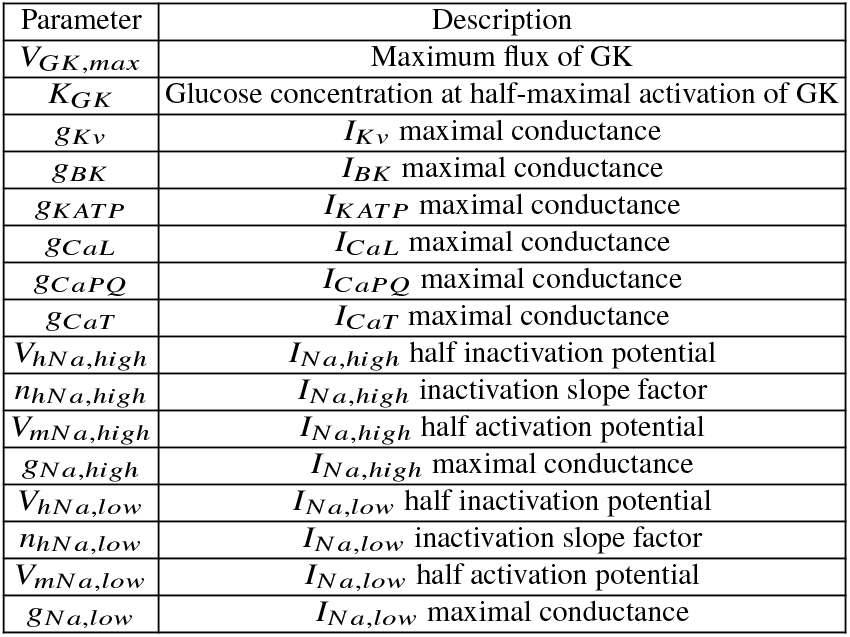
Heterogeneous parameters in the *β*-cells ODE model.

| Parameter | Description |
| --- | --- |
| $V_{GK,max}$ | Maximum flux of GK |
| $K_{GK}$ | Glucose concentration at half-maximal activation of GK |
| $g_{Kv}$ | $I_{Kv}$ maximal conductance |
| $g_{BK}$ | $I_{BK}$ maximal conductance |
| $g_{KATP}$ | $I_{KATP}$ maximal conductance |
| $g_{CaL}$ | $I_{CaL}$ maximal conductance |
| $g_{CaPQ}$ | $I_{CaPQ}$ maximal conductance |
| $g_{CaT}$ | $I_{CaT}$ maximal conductance |
| $V_{hNa,high}$ | $I_{Na,high}$ half inactivation potential |
| $n_{hNa,high}$ | $I_{Na,high}$ inactivation slope factor |
| $V_{mNa,high}$ | $I_{Na,high}$ half activation potential |
| $g_{Na,high}$ | $I_{Na,high}$ maximal conductance |
| $V_{hNa,low}$ | $I_{Na,low}$ half inactivation potential |
| $n_{hNa,low}$ | $I_{Na,low}$ inactivation slope factor |
| $V_{mNa,low}$ | $I_{Na,low}$ half activation potential |
| $g_{Na,low}$ | $I_{Na,low}$ maximal conductance |

Using this framework, we implement a categorical feature-based sensitivity analysis to quantify how the heterogeneous parameters influence different *β*-cell phenotypes (or features). The dynamic behaviors of the *β*-cells are classified into *n*_*c*_ = 5 phenotypes: silent, depolarized, spiking, bursting, and other. These phenotypes are determined from the raw membrane potential time-series by the algorithm in Figure 5.3. We applied logistic regression to binarized classification of each phenotype and standardized parameter deviations. The associated input matrix 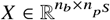 contains the standardized parameter values for each of the *n*_*b*_ cells to account for differences in scale before fitting the logistic regression model.

**Fig. 5.3:**
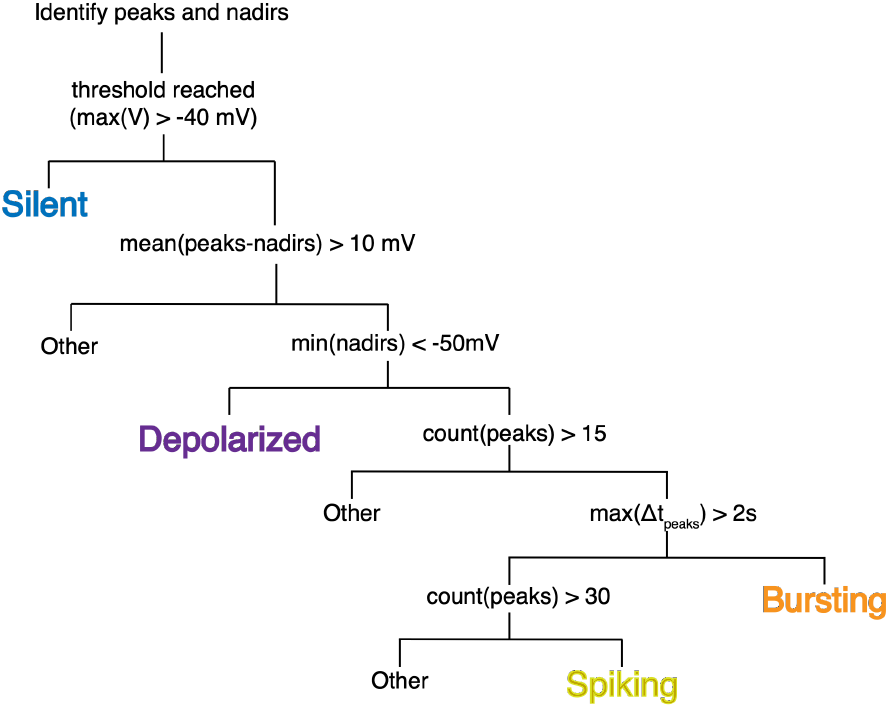
A phenotype classification algorithm used to determine the dynamic behavior of *β*-cells in response to glucose. The algorithm depends on the simulated voltage behavior in terms of its highest points (peaks), lowest points (nadirs), and peak-to-peak interval (Δ*t*_peaks_). More than 1000 automatic classifications were randomly selected and verified for correctness by the same analyst. This algorithm is very similar to the one developed in [27] on a heterogeneous population built from the same dataset.

**Fig. 5.4:**
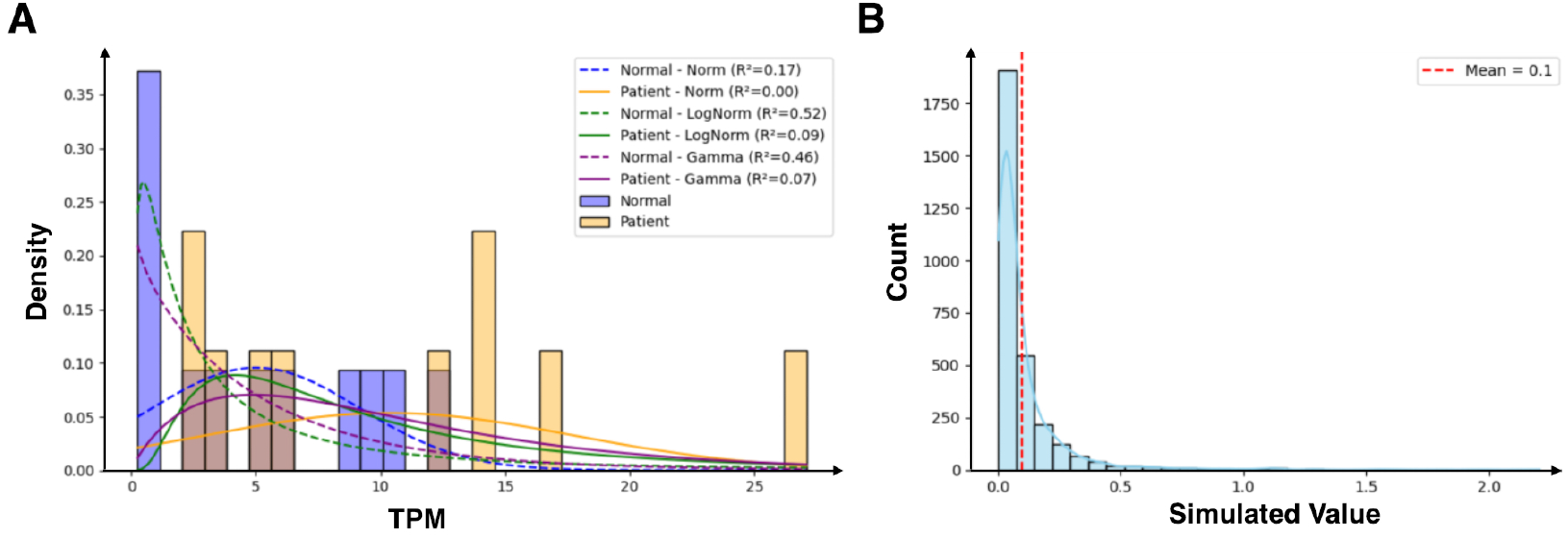
Distribution of KCNN3 expression from RNA-seq and parameterization of *g*_*SK*_. (A) Distribution fitting of *KCNN3* transcript expression. Normal, log-normal, and gamma distributions were fitted to TPM values of *KCNN3* in *β*-cells. In healthy donors, expression was best captured by a log-normal distribution (*R*^2^ ≈ 0.52) (B) Parameterization of *g*_*SK*_ from the fitted distribution. *g*_*SK*_ was parameterized using values sampled from the log-normal distribution of *KCNN3* transcript expression in healthy *β*-cells, with the mean aligned to 0.1 nS/pF. All simulations were run for 90 s.

### 5.2.2 Transcriptomic analysis and modeling approach

We reanalyzed the single-cell RNA sequencing dataset in [20], which profiled over 2,000 pancreatic islet cells from healthy donors and patients with type 2 diabetes. After filtering for *β*-cells, we performed separate analyses for healthy and diabetic groups, quantifying KCNN3 transcript expression as a proxy for SK channel abundance. Normal, lognormal, and gamma distributions were fitted to transcript-per-million (TPM) values of KCNN3. In *β*-cells from healthy donors, expression followed a skewed distribution that was best captured by a log-normal model (r^2^ ≈ 0.52). In contrast, *β*-cells from patients with type 2 diabetes showed lower expression and greater variability, and none of the tested models provided a satisfactory fit. For simulations, SK conductance (*g*_*SK*_) was parameterized using values sampled from the log-normal distribution of KCNN3 transcript expression observed in healthy *β*-cells, with the mean set to 0.1 nS/pF. The SK channel current was modeled using a standard Hill-type formulation:

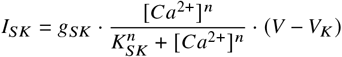

In this equation, *g*_*SK*_ denotes the maximal SK conductance, *Ca*^2+^ represents the intracellular calcium concentration, *K*_*SK*_ is the half-activation constant, and *n* is the Hill coefficient describing the cooperativity of calcium binding. The term *V V*_*K*_ corresponds to the driving force for potassium ions, with *V*_*K*_ being the potassium reversal potential. This formulation captures the calcium-dependent activation of SK channels and their hyperpolarizing influence on the membrane potential. Additional simulations were conducted with fixed *g*_*SK*_ values of 0.001, 0.01, 0.1, and 1 nS/pF for comparison. The value of 0.1 nS/pF served as the reference, corresponding to the conductance used in prior modeling studies [26] based on experimental data. All simulations, both fixed and heterogeneous, were run for 90 s. Cell phenotypes were classified according to the same criteria described in the previous section.

### 5.2.3 α-β-Cell communication

#### 5.2.3.1 Updated Cha-Noma and Coupled Cha-Noma Model

A previously published mathematical model describing a single mouse *β*-cell [28], termed the Cha-Noma model, as well as an extension of that model to an islet [16], termed the coupled Cha-Noma model, was used as a starting point. Both describe the dynamic activity of *β*-cell membrane potential, *V*_*m*_, as a function of 8 plasma membrane currents and 3 ion transporters. The time-dependent change in voltage for the Cha-Noma model is written as follows:

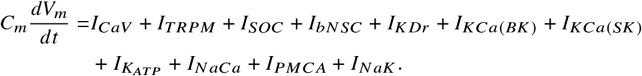

The coupled Cha-Noma model extends this model with the introduction of a coupling current to the right-hand side of the above equation,

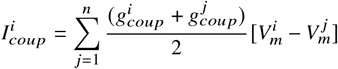

as well as the introduction of heterogeneity in several cellular parameters. This heterogeneity is the same as described in [29]. Both the single cell and couple Cha-Noma models were updated to include the effect of *α*-*β*-cell communication on ATP-sensitive K^+^channels, by introducing a GLP-1R signaling model described in [30]. This allowed GLP-1 concentration to stimulate production of intracellular cyclic-AMP and, further, Protein Kinase A activity. The effects of activated PKA (PKA_*a*_) on K_ATP_ channels were modeled using a formulation from [31] in which the available MgADP to alter the K_ATP_ channel open probability decreases as PKA_*a*_ increases, shown as follows:

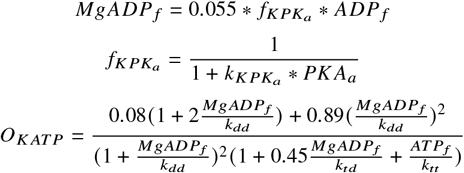

The open probability then affects the K_ATP_ channel current as follows:

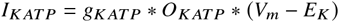

This resulted in an updated model, shown schematically in Figure 5.5.

**Fig. 5.5:**
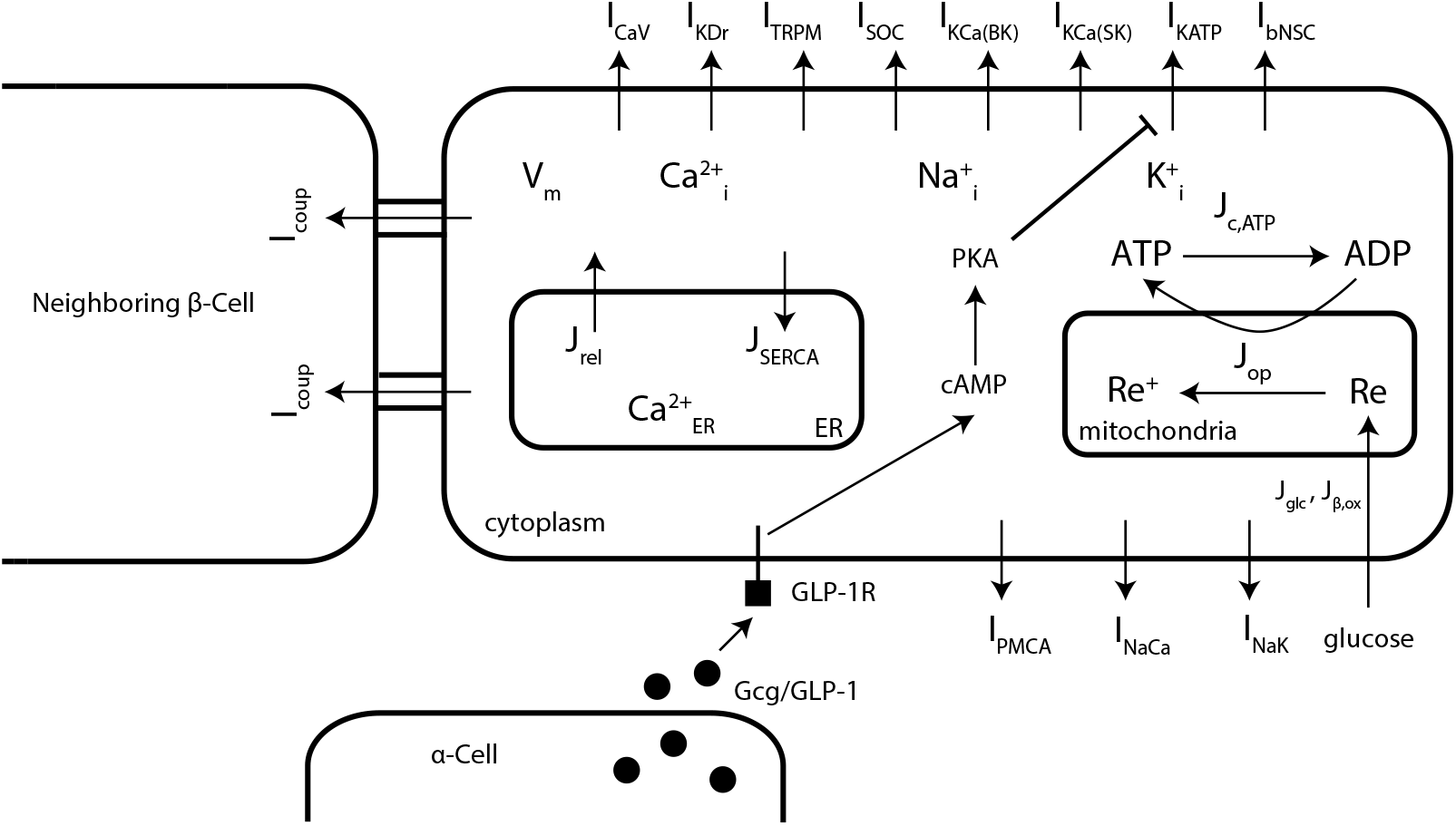
Updated coupled Cha-Noma model, modified from [28]. This updated model incorporates *α*-*β*-cell communication via GLP-1 acting on the GLP-1R, resulting in an inhibitory effect on *K*_*ATP*_ channels. The updated single-cell Cha Noma model is similar, without the inclusion of *β*-*β*-cell coupling.

To introduce *α*-*β*-cell communication into the coupled Cha-Noma model, it was also necessary to alter the islet structure. This was necessary as the previous iteration of the coupled Cha-Noma model only included *β*-cells, but the introduction of *α*-*β*-cell communication requires two things: 1) the locations of *α*-cells, and 2) which *β*-cells are located close enough to an *α*-cell to be affected by paracrine signaling. Both of these requirements were met by applying a modified version of the simulated annealing algorithm developed previously by Felíx-Martinez et al. [14, 15]. This allowed the reconstruction of a mouse islet from published immunofluorescence data that included the cell locations for both *α*-cells and *β*-cells [32]. To determine cell contacts, a threshold between two cell surfaces of 6 *μm* was used. This threshold was chosen for the fact that it gave a similar *β*-*β* link distribution to that in the coupled Cha-Noma model, allowing for comparisons between the two versions of the models if so desired. The updated islet structure and *β*-*β* structural degree distributions can be seen in Figure 5.6.

**Fig. 5.6:**
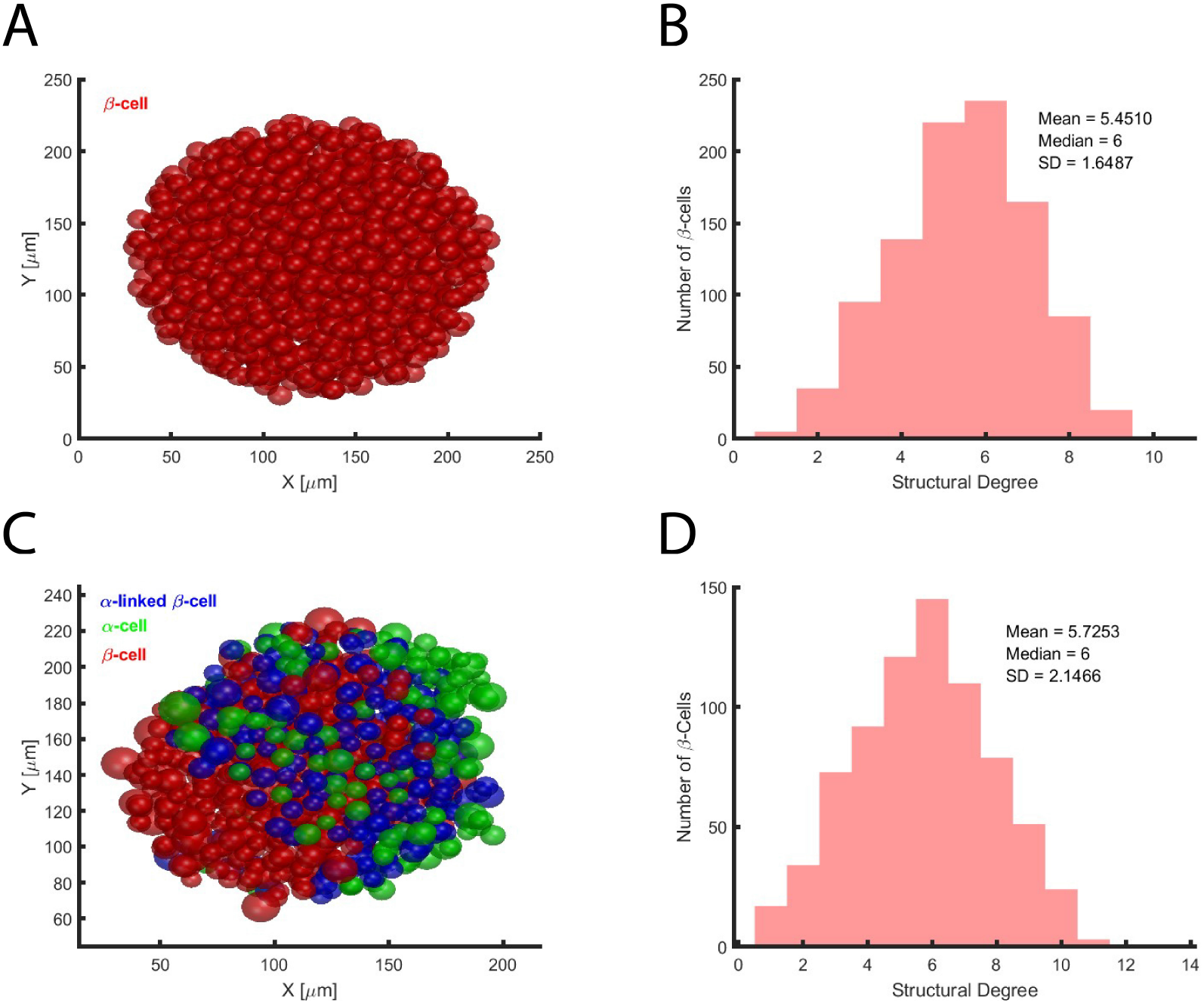
Updated islet structure. (A) Islet structure in the coupled Cha-Noma model. (B) *β*-*β* structural distribution for the coupled Cha-Noma model. (C) Updated islet structure that includes *α*-cells. (D) *β*-*β* structural distribution for the updated version of the coupled Cha-Noma model

#### 5.2.3.2 Cha-Noma Model Simulations

Simulations were run using both the updated single-cell Cha-Noma model and the updated coupled Cha-Noma model. The parameters in the single-cell model were set to the mean of those in the coupled Cha-Noma model. For both models, the applied glucose followed a step function, in which glucose was raised from 2 mM to 11 mM at 20 s. For simulations with the coupled model, two sets were run. In the first, [GLP-1] was set to 0 nM across all *β*-cells. This was done to represent no *α*-*β*-cell communication. In the second, [GLP-1] was set to 20 nM only for *α*-linked *β*-cells. This was done to represent an islet with *α*-*β*-cell communication. A concentration of 20 nM was chosen as it produces maximal stimulation of the GLP-1R. A total of 6 seeds with randomized cellular heterogeneity were used while maintaining the same islet structure.

#### 5.2.3.3 First Responder and Time of Response Analysis

For the multicellular model, the time of response for individual *β*-cells was calculated using individual calcium traces and a custom MATLAB script. Time of response was calculated as the point at which [Ca^2+^] reached 25% of its maximum value relative to the steady state [Ca^2+^] during 2 mM. This time of response was then made relative to the time at which glucose was elevated by subtracting 20 s. A threshold of 25 % was chosen as it gave reasonable results upon visual inspection. First responders were set as the 75 fastest responding cells (out of 750 total). Some examples of the points chosen using this method can be seen in Figure 5.7. For the single-cell model, time of response could be determined by hand.

**Fig. 5.7:**
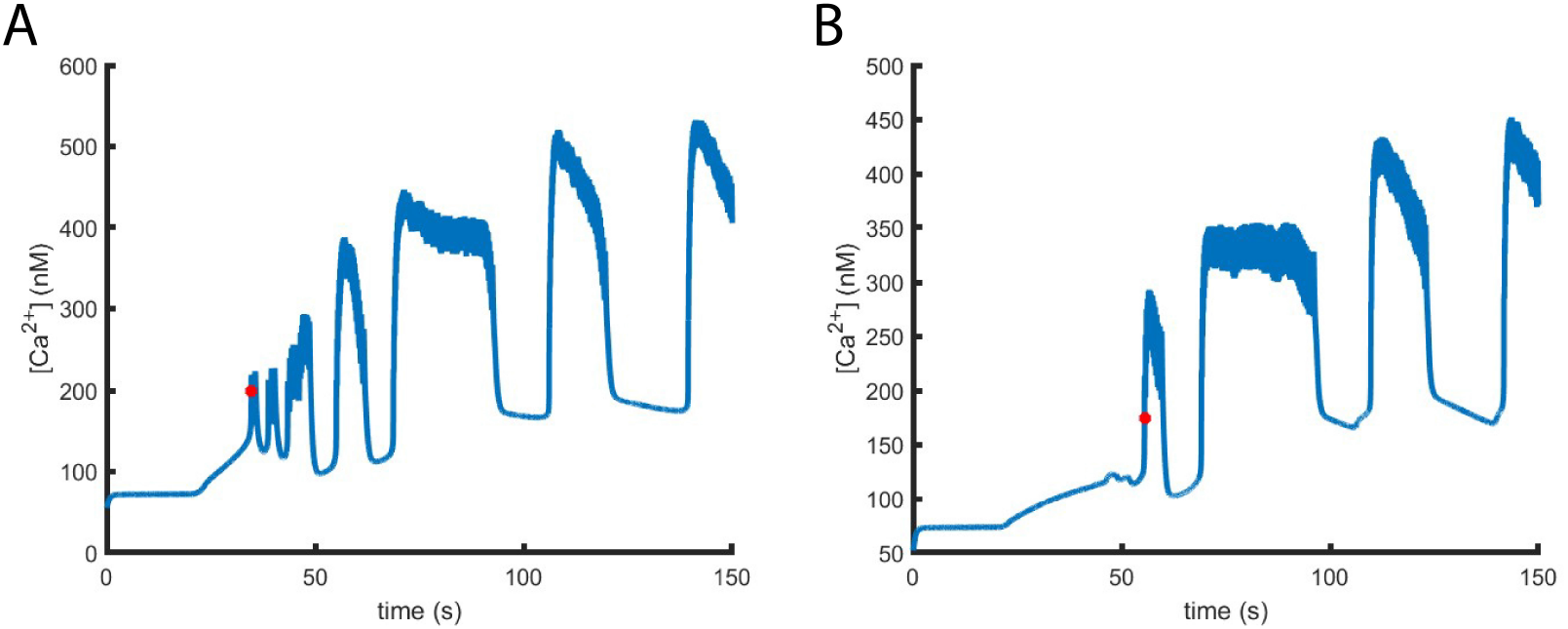
Example time of response points. (A) Example cell in which the point the islet is presumed to be responding is indicated in red. (B) Example cell in which the point the islet is presumed to be responding is indicated in red.

## 5.3 Results

### 5.3.1 Sensitivity Analysis of Riz Model

As mentioned in the methods, we simulated the Riz (human) *β*-cell population model at a glucose concentration of 20 mM for 600 s. We the phenotype classification algorithm in Figure 5.3 and determine the frequency of the resulting phenotypes as shown in Figure 5.8 below.

**Fig. 5.8:**
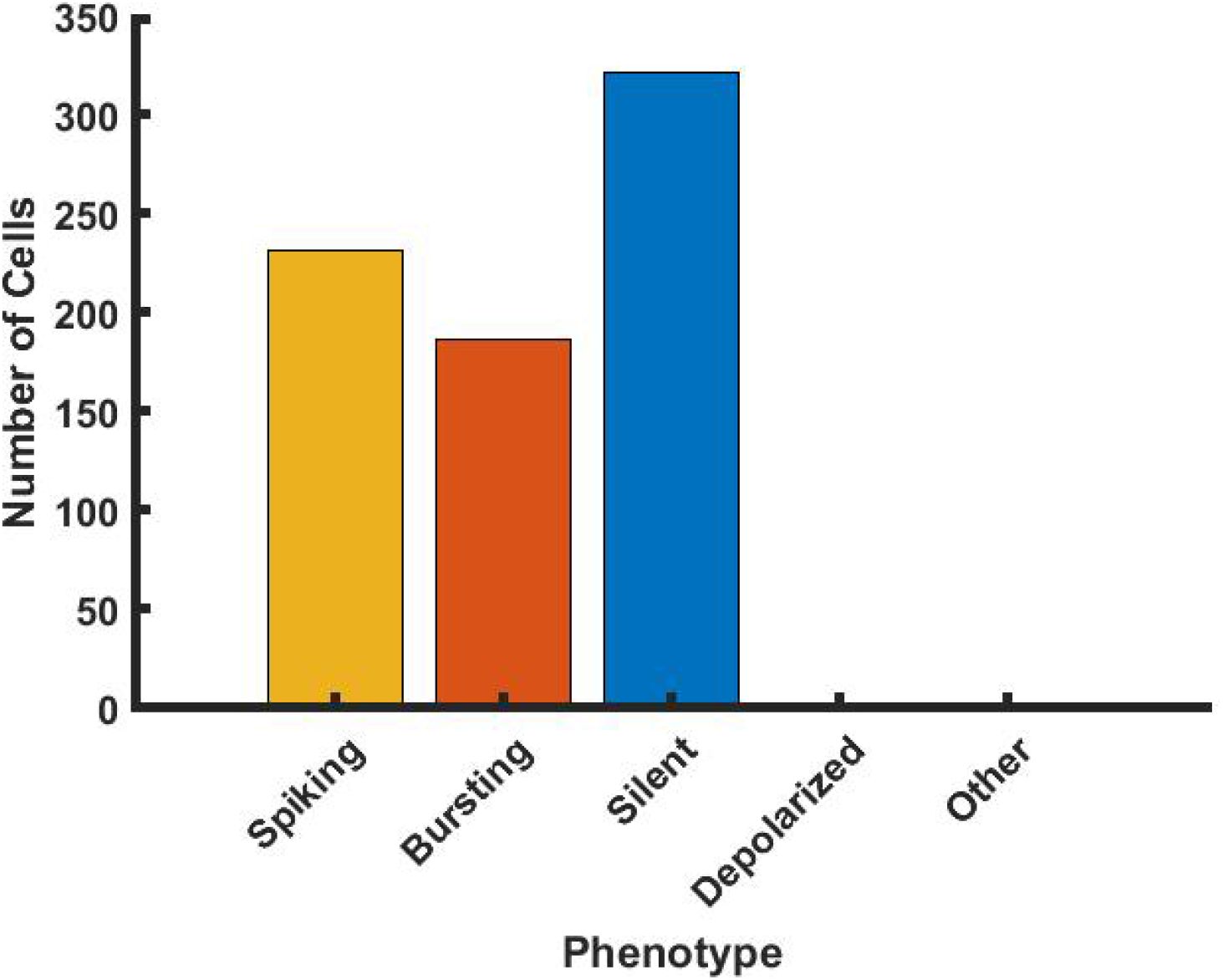
Occurrence of phenotypes of *β*-cells in Riz model in the case of 20 mM glucose.

The predominant phenotypes are spiking, bursting, and silent, with the remaining phenotypes (depolarized and other) accounting for *<* 1% of cells. Thus, we disregard the other two phenotypes for our logistic regression sensitivity analysis because those lesser phenotypes would be dramatically underpowered and determinant parameters would not be reliably identifiable. Recall from the methods that the logistic regression approach used has an input matrix ∈ *X* R^744×16^ with 744 *β*-cells and 16 parameters. This matrix includes two groups of parameters that determine whether the Na^+^ current exhibits high or low steady-state voltage dependence. As a result, each cell contains 4 columns in which 1 group of Na^+^ current parameters that are zero (because that form of the current is not expressed). This results in a rank-deficient structure, which leads to an ill-conditioned logistic regression problem. To address this, we reduce the dimension of the input matrix *X* by consolidating the nonzero entries of Na^+^ current parameters, resulting in a simplified matrix *X*^*S*^ ∈ ℝ^744×12^. Now, we apply the logistic regression approach with the updated input matrix *X* ^*S*^ to retrieve a regression coefficient matrix containing the sensitivities of the different parameters for each phenotype. These are plotted in 5.9 below.

Figure 5.9 reveals that *g*_*K ATP*_, *g*_*BK*_, and *V*_*hN a*_ are the most sensitive parameters in the case of both spiking and silent phenotypes and *K*_*GK*_, *g*_*BK*_, and *V*_*GK*,max_ are the most sensitive parameters for the bursting phenotype. Since we are interested in exaggerating the bursting phenotype, we modulated the variability of the most sensitive parameters for the bursting phenotype by multiplying the mean of the distribution that generates these parameter values by half for parameters *K*_*GK*_ and *g*_*BK*_ and the mean of the distribution governing *V*_*GK*,max_ by 2. We run the simulation of the human population again, generating the new phenotypes and then assess the change in occurrence of each phenotype at the same high glucose concentration (20 *mM*). This change is reflected in Figure 5.10 below.

**Fig. 5.9:**
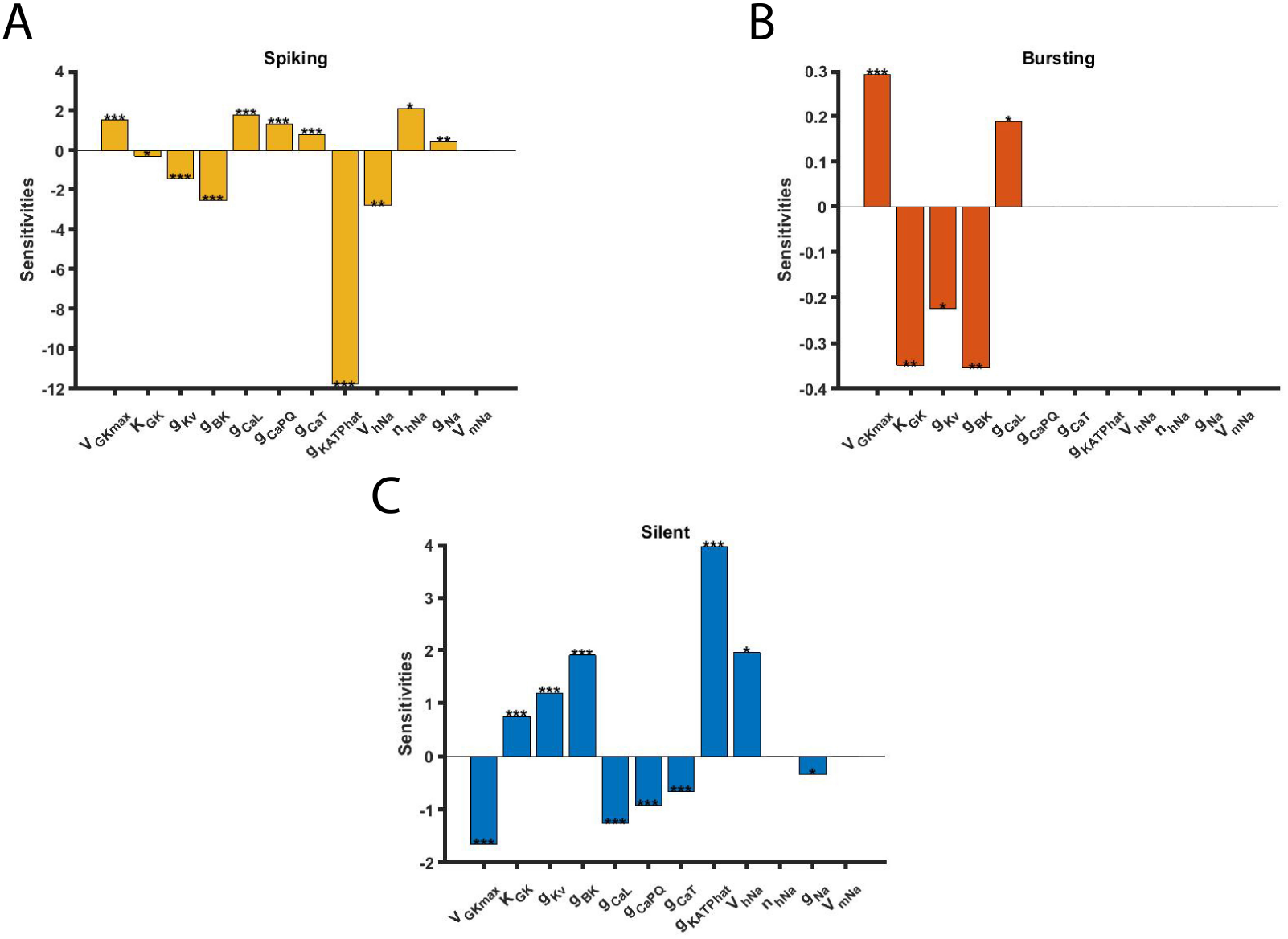
Sensitivity analysis. (A) Parameter sensitivities for spiking phenotype (B) Parameter sensitivities for bursting phenotype (C) Parameter sensitivities for silent phenotype. (*p-value¡0.05, **p-value¡0.01, and ***p-value¡0.001).

**Fig. 5.10:**
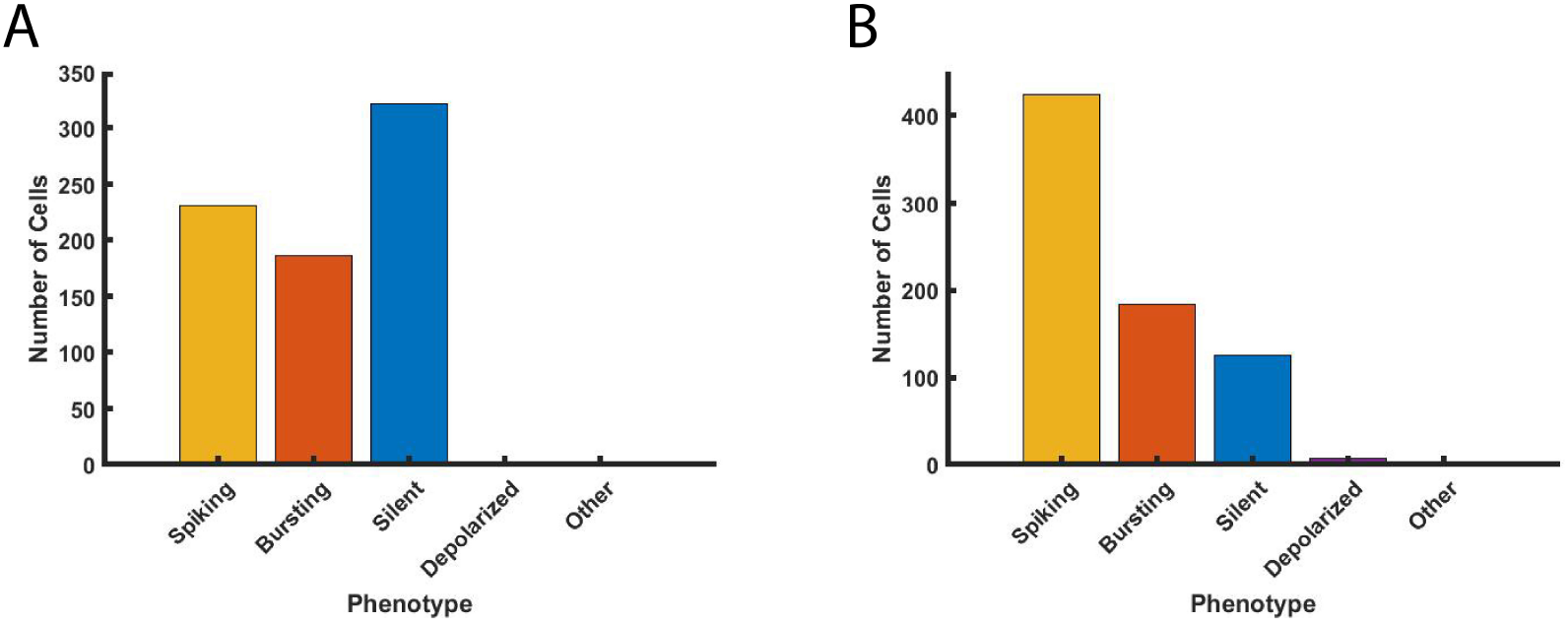
Occurrence of phenotypes of *β*-cells in Riz model in the case of 20 mM glucose. (A) before changing the variability of the most sensitive parameters for the bursting phenotype (B) after changing the variability of the most sensitive parameters (*K*_*GK*_, *g*_*BK*_, and *V*_*GK*,max_) for the bursting phenotype.

**Fig. 5.11:**
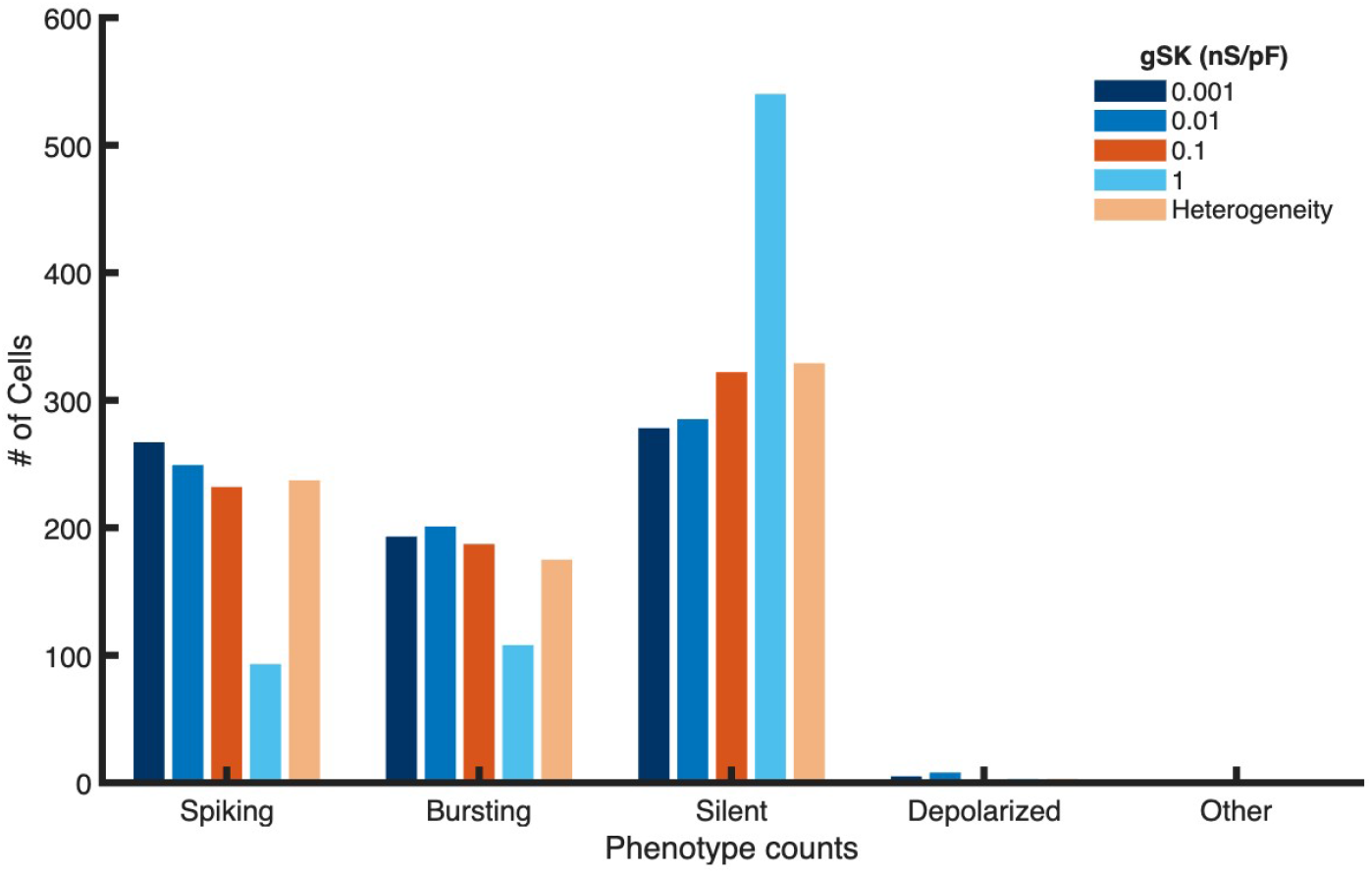
Comparison of *β*-cell phenotype analysis under fixed and heterogeneous SK conductance. Phenotypes were classified as spiking, bursting, silent, depolarized, or other. Fixed-conductance groups were simulated at 0.001, 0.01, 0.1 (reference), and 1 nS/pF. In addition, a heterogeneous group was generated by sampling *g*_*SK*_ values from the log-normal distribution of *KCNN3* transcript expression obtained from RNA-seq data, with the mean aligned to 0.1 nS/pF. All simulations were run for 90 s.

**Fig. 5.12:**
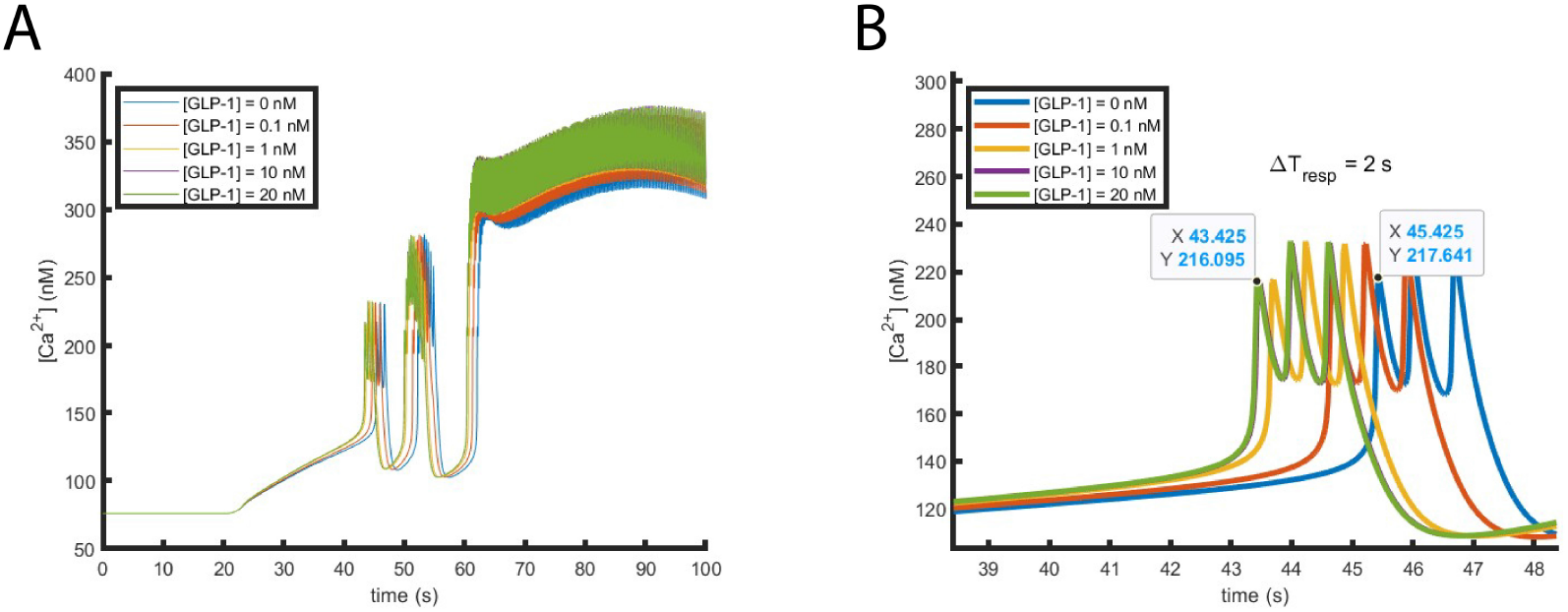
Effects of GLP-1 stimulation on single-cell model. (A) Effects of 0, 0.1, 1, 10, and 20 nM GLP-1 stimulation on the 1^st^ phase response. (B) Zoomed in version of A, showing a maximal leftward shift in time of response of 2 s.

Counter to our expectations, this approach exaggerated the prevalence of the spiking phenotype in the modified population, rather than enhancing the prevalence of bursting. This is because, even though *K*_*GK*_, *g*_*BK*_, and *V*_*GK*,max_ are the most sensitive determinants of bursting, they are also sensitive modulators of spiking, and reciprocally so 5.9. As a result, they are not independently able to exaggerate the bursting phenotype within the population. Thus, we next investigated the effect of targeting the variability of the SK current in the Riz model, which, as mentioned above, was previously implemented as invariant within the *β*-cell population.

### 5.3.2 Comparison of *β*-cells phenotypes under fixed and heterogeneous SK conductance

To investigate how variability in SK channel conductance influences *β*-cells electrical behavior, we compared cell phenotypes generated under fixed SK conductance values (0.001, 0.01, 0.1, and 1 nS/pF) with those observed in heterogeneous populations sampled from the log-normal distribution of KCNN3 transcript expression. While distribution fitting confirmed that KCNN3 expression in *β*-cells follows a log-normal pattern in healthy donors, the central question was whether such heterogeneity in SK conductance would affect the diversity of electrophysiological outcomes in our model. Simulations with fixed SK conductance values revealed distinct trends across groups. Both lower-conductance groups (0.001 and 0.01 nS/pF) exhibited more bursting and spiking than the reference value of 0.1 nS/pF. Within these, the 0.01 nS/pF group showed the highest proportion of bursting activity but reduced spiking compared with the 0.001 nS/pF group. In contrast, the 0.1 nS/pF group displayed fewer active cells overall, with reduced spiking and bursting but a higher proportion of silent states than either of the lower-conductance groups. At the highest conductance of 1 nS/pF, most cells became silent, reflecting excessive repolarizing drive that strongly suppressed excitability. When SK conductance values were sampled from the log-normal distribution of KCNN3 transcript expression, the simulated population exhibited a mixture of spiking, bursting, and silent cells. The overall distribution of phenotypes was similar to that observed at the reference value of 0.1 nS/pF, which has been used in previous models based on experimental data. This similarity indicates that incorporating heterogeneity does not disrupt the established parameterization of the model or require compensatory adjustments of other parameters. Instead, the introduction of a heterogeneous population reflects the physiological reality that SK channel expression varies across *β*-cells, while maintaining compatibility with the existing model framework. Thus, the key contribution of this approach is that it updates the model toward a more faithful representation of biological diversity without sacrificing internal consistency.

### 5.3.3 GLP-1 mediated inhibition of *β*-cell *I*_*K AT P*_ plays a subtle but measurable role in promoting 1st responder behavior

As mentioned in the methods, *α*-*β*-cell communication was introduced into both the Cha-Noma model and the coupled Cha-Noma model. The effect was modeled as an inhibition of the K_ATP_ channel. To first get an idea of how the average cell would be affected by *α*-*β*-cell communication, the single-cell model was run with various applied concentrations of GLP-1 (0, 0.1, 1, 10, 20) nM. At maximal stimulation (20 nM GLP-1), the calcium curve was leftward shifted by 2 s. This is not a large effect, and likely to be at the border of detectability in analogous experiments, but given that it only involves one of many known influences of GLP-1 signaling on *β*-cell excitability, it suggests that this mode of *α*-*β*-cell communication may be quantitatively important in the first phase glucose response.

Next, the effect of *α*-*β*-cell communication on the 1^st^ responders subpopulation was quantified by running 6 seeds of the multicellular model for two separate conditions as described in the methods. Time of response and 1^st^ responders were determined for both control (0 nM GLP-1) and stimulatory (20 nM GLP-1) experiments. Then, the number of 1^st^ responders that were *α*-linked in both control and stimulatory conditions were calculated and compared using a Wilcoxon signed rank test. The results are shown in Figure 5.13.

**Fig. 5.13:**
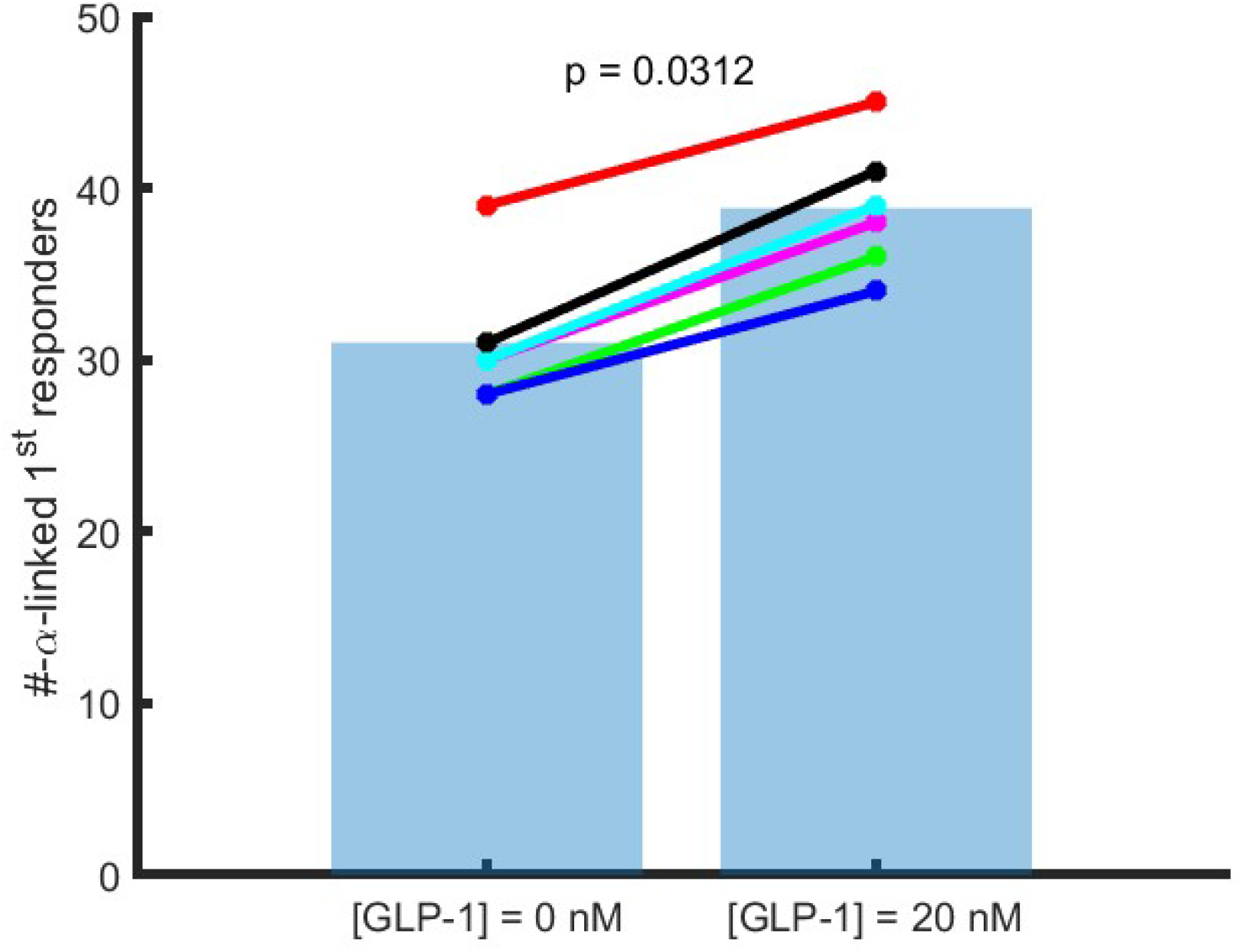
Effect of *α*-*β*-cell communication on 1^st^ responder subpopulation. Statistical signficance testing done using a Wilcoxon signed rank test.

From Figure 5.13, it can be seen that introducing *α*-*β*-cell communication increased the number of *α*-linked *β*-cells that were 1^st^ responders. This trend was observed across all six seeds.

## 5.4 Discussion

In this manuscript, we aim to reconstruct realistic islets by studying the effect of heterogeneous parameters and some constant parameters of *β*-cells as well as paracrine signaling for *α*-*β*-cell communication on intrinsic and extrinsic heterogeneity. One of the ways to reach a successful reconstruction was to change the number of bursting cells to help coordinate glucose-dependent activity.

While the sensitivity analysis approach used above guided our search for the most sensitive parameters for the bursting *β*-cell phenotype, the parameter sensitivities for spiking cells are dominant, making it difficult to selectively modulate the bursting phenotype without concurrently influencing the spiking and silent populations. These findings highlight the complexity of phenotype regulation within the *β*-cell population and point to the need for more targeted multi-parameter adjustments or nonlinear sensitivity analyses to independently modulate the bursting phenotype. It also supports the contention that the crucial bursting dynamics may in fact not be a common property of single *β*-cells but rather, as has been argued previously [33], an emergent dynamic resulting from coupling cells with silent and spiking phenotypes.

Our comparison of *β*-cell phenotypes under fixed versus heterogeneous SK conductance demonstrates that variability in SK channel expression can be incorporated into computational models without disrupting established behavior. At lower conductance levels (0.001 and 0.01 nS/pF), spiking and bursting were more prevalent than at the reference value of 0.1 nS/pF. In contrast, at the highest conductance (1 nS/pF), nearly all cells became silent due to excessive repolarizing drive. Under the heterogeneous condition, the resulting phenotype distribution closely resembled that of the reference group (0.1 nS/pF). This similarity indicates that the introduction of heterogeneity does not compromise the internal consistency of the model or necessitate compensatory adjustments of other parameters. Instead, it allows the model to retain its established dynamics while explicitly reflecting the biological fact that SK channel expression is heterogeneous among *β*-cells. In this sense, the major contribution of our approach lies not in altering the predicted outcomes relative to the reference conductance, but in updating the framework to incorporate physiologically grounded variability.

From a biological perspective, embedding heterogeneity at the channel level provides a more faithful representation of *β*-cell networks, where functional diversity among cells is an experimentally observed hallmark. From a modeling perspective, this approach increases the fidelity of simulations by aligning them more closely with the molecular data, thereby providing a stronger foundation for studying both normal physiology and disease-related dysfunction. In particular, because *β*-cell heterogeneity is disrupted in type 2 diabetes, models that integrate transcriptomic variability may offer valuable tools for probing how such changes contribute to impaired islet function.

Introduction of *α*-*β*-cell communication via GLP-1 acting on the GLP-1R to decrease the K_ATP_ channel conductance resulted in a statistically significant increase in *α*-linked *β*-cells being defined as 1^st^ responders. However, this role seems to be minor as indicated by the number of *α*-linked *β*-cells that were already 1^st^ responders prior to GLP-1 stimulation. This indicates that intrinsic parameters and gap-junction coupling are likely to also play substantial roles in defining the 1^st^ subpopulation, as shown in previous work [29].

It is also important to address some limitations of this implementation of *α*-*β*-cell communication. The first being that only one effect of GLP-1R signaling was modeled. As detailed in this review [34], the effects of increased cyclic-AMP production on *β*-cells function are various. Thus, future models must work to incorporate these effects to get a true understanding of how paracrine signaling between *α* and *β*-cells impacts *β*-cells function. Second, PKA_*a*_ was modeled as being an instantaneous function of cyclic-AMP. However, previous studies have indicated that PKA_*a*_ shows a distinct relationship to cyclic-AMP oscillations [35]. Third, the strength and distance of paracrine signaling within the islet is not known. In this model, only *β*-cells that were in contact with *α*-cells were considered to be close enough for paracrine interactions. However, in real islets this may not be the case. Furthermore, the actual extracellular concentration of GLP-1 or glucagon that an *α*-linked *β*-cell might be exposed to is unknown, but, for simplicity, was set to a concentration that produces maximal stimulation. Fourth, calculating the time of response with the given population of *β*-cells proved to be difficult owing to the spiking nature of their first phase. A simple method of the time point at which the [Ca^2+^] reached 25 % of its maximum value relative to the steady state [Ca^2+^] at 2 mM glucose was used.

A separate method that utilized MATLABs findpeaks() function was also employed, and gave a similar, statistically significant result. Manual inspection of the time points chosen for both methods indicated them to be reasonable, but not perfect. Thus, a more rigorous method should be employed in future work to confirm the results.

## References

1. Over Cabrera, Dora M. Berman, Norma S. Kenyon, Camillo Ricordi, Per-Olof Berggren, and Alejandro Caicedo. The unique cytoarchitecture of human pancreatic islets has implications for islet cell function. Proceedings of the National Academy of Sciences, 103(7):2334–2339, 2006.

2. Danh-Tai Hoang, Hitomi Matsunari, Masaki Nagaya, Hiroshi Nagashima, J. Michael Millis, Piotr Witkowski, Vipul Periwal, Manami Hara, and Junghyo Jo. A conserved rule for pancreatic islet organization. PLOS ONE, 9(10):1–9, 10 2014.

3. German Kilimnik, Junghyo Jo, Vipul Periwal, Mark C. Zielinski, and Manami Hara. Quantification of islet size and architecture. Islets, 4(2):167–172, 2012. PMID: 22653677.

4. Abraham Kim, Kevin Miller, Junghyo Jo, German Kilimnik, Pawel Wojcik, and Manami Hara. Islet architecture: A comparative study. Islets, 1(2):129–136, 2009. PMID: 20606719.

5. Marcela Brissova, Michael J. Fowler, Wendell E. Nicholson, Anita Chu, Boaz Hirshberg, David M. Harlan, and Alvin C. Powers. Assessment of human pancreatic islet architecture and composition by laser scanning confocal microscopy. Journal of Histochemistry & Cytochemistry, 53(9):1087–1097, 2005. PMID: 15923354.

6. Joan Camunas-Soler, Xiao-Qing Dai, Yan Hang, Austin Bautista, James Lyon, Kunimasa Suzuki, Seung K. Kim, Stephen R. Quake, and Patrick E. MacDonald. Patch-Seq Links Single-Cell Transcriptomes to Human Islet Dysfunction in Diabetes. Cell Metabolism, 31(5):1017–1031.e4, May 2020. Publisher: Elsevier.

7. TL Jetton and MA Magnuson. Heterogeneous expression of glucokinase among pancreatic beta cells. Proceedings of the National Academy of Sciences, 89(7):2619–2623, April 1992. Publisher: Proceedings of the National Academy of Sciences.

8. R. Kiekens, P. In ‘t Veld, T. Mahler, F. Schuit, M. Van De Winkel, and D. Pipeleers. Differences in glucose recognition by individual rat pancreatic B cells are associated with intercellular differences in glucose-induced biosynthetic activity. The Journal of Clinical Investigation, 89(1):117–125, January 1992. Publisher: The American Society for Clinical Investigation.

9. David W. Piston, Susan M. Knobel, Catherine Postic, Kathy D. Shelton, and Mark A. Magnuson. Adenovirus-mediated Knockout of a Conditional Glucokinase Gene in Isolated Pancreatic Islets Reveals an Essential Role for Proximal Metabolic Coupling Events in Glucose-stimulated Insulin Secretion *. Journal of Biological Chemistry, 274(2):1000–1004, January 1999. Publisher: Elsevier.

10. M. Van De Winkel and D. Pipeleers. Autofluorescence-activated cell sorting of pancreatic islet cells: Purification of insulin-containing B-cells according to glucose-induced changes in cellular redox state. Biochemical and Biophysical Research Communications, 114(2):835–842, July 1983.

11. D. Mears, N. F. Sheppard, I. Atwater, and E. Rojas. Magnitude and modulation of pancreatic β-cell gap junction electrical conductance in situ. J. Membarin Biol., 146(2):163–176, July 1995.

12. M. Pérez-Armendariz, C. Roy, D. C. Spray, and M. V. Bennett. Biophysical properties of gap junctions between freshly dispersed pairs of mouse pancreatic beta cells. Biophysical Journal, 59(1):76–92, January 1991.

13. Nikki L. Farnsworth, Alireza Hemmati, Marina Pozzoli, and Richard K. P. Benninger. Fluorescence recovery after photobleaching reveals regulation and distribution of connexin36 gap junction coupling within mouse islets of Langerhans. The Journal of Physiology, 592(20):4431–4446, 2014. eprint: https://physoc.onlinelibrary.wiley.com/doi/pdf/10.1113/jphysiol.2014.276733.

14. Gerardo J. Félix-Martínez, Aurelio N. Mata, and J. Rafael Godínez-Fernández. Reconstructing human pancreatic islet architectures using computational optimization. Islets, 12(6):121–133, November 2020. Publisher: Taylor & Francis eprint: 10.1080/19382014.2020.1823178.

15. Gerardo J. Félix-Martínez and J.R. Godínez-Fernández. Comparative analysis of reconstructed architectures from mice and human islets. Islets, 14(1):23–35, December 2022. Publisher: Taylor & Francis eprint: 10.1080/19382014.2021.1987827.

16. JaeAnn M. Dwulet, Nurin W. F. Ludin, Robert A. Piscopio, Wolfgang E. Schleicher, Ong Moua, Matthew J. Westacott, and Richard K. P. Benninger. How Heterogeneity in Glucokinase and Gap-Junction Coupling Determines the Islet [Ca2+] Response. Biophysical Journal, 117(11):2188–2203, December 2019. Publisher: Elsevier.

17. Roshni Shetty, Radhika Singh-Agarwal, Stefan Meier, Christian Goetz, and Andrew G. Edwards. Reconstruction of a pancreatic beta cell network from heterogeneous functional measurements. In Computational Physiology: Simula Summer School 2023 - Student Reports, pages 71–86. Springer Nature Switzerland, Cham, 2024.

18. Karoline Horgmo Jæger and Aslak Tveito. Sometimes extracellular recordings fail for good reasons. bioRxiv, July 2025. ISSN: 2692-8205 Pages: 2025.07.01.662690 Section: New Results.

19. Reshma Ramracheya, Caroline Chapman, Margarita Chibalina, Haiqiang Dou, Caroline Miranda, Alejandro González, Yusuke Moritoh, Makoto Shigeto, Quan Zhang, Matthias Braun, et al. Glp-1 suppresses glucagon secretion in human pancreatic alpha-cells by inhibition of p/q-type ca2+ channels. Physiological reports, 6(17):e13852, 2018.

20. Åsa Segerstolpe, Athanasia Palasantza, Pernilla Eliasson, Eva-Marie Andersson, Anne-Christine Andréasson, Xiaoyan Sun, Simone Picelli, Alan Sabirsh, Maryam Clausen, Magnus K Bjursell, et al. Single-cell transcriptome profiling of human pancreatic islets in health and type 2 diabetes. Cell metabolism, 24(4):593–607, 2016.

21. Adam P. Chambers, Joyce E. Sorrell, April Haller, Karen Roelofs, Chelsea R. Hutch, Ki-Suk Kim, Ruth Gutierrez-Aguilar, Bailing Li, Daniel J. Drucker, David A. D’Alessio, Randy J. Seeley, and Darleen A. Sandoval. The Role of Pancreatic Preproglucagon in Glucose Homeostasis in Mice. Cell Metab, 25(4):927–934.e3, April 2017.

22. Over Cabrera, James Ficorilli, Janice Shaw, Felipe Echeverri, Frank Schwede, Oleg G. Chepurny, Colin A. Leech, and George G. Holz. Intra-islet glucagon confers β-cell glucose competence for first-phase insulin secretion and favors GLP-1R stimulation by exogenous glucagon. Journal of Biological Chemistry, 298(2):101484, February 2022.

23. Megan E. Capozzi, Berit Svendsen, Sara E. Encisco, Sophie L. Lewandowski, Mackenzie D. Martin, Haopeng Lin, Justin L. Jaffe, Reilly W. Coch, Jonathan M. Haldeman, Patrick E. MacDonald, Matthew J. Merrins, David A. D’Alessio, and Jonathan E. Campbell. β Cell tone is defined by proglucagon peptides through cAMP signaling. JCI Insight, 4(5), March 2019. Publisher: American Society for Clinical Investigation.

24. Nirmala V. Balasenthilkumaran, Lidija Križančić Bombek David Ramirez, Maša Skelin Klemen, Eva Paradiž Leitgeb, Jasmina Kerčmar, Jan Kopecky, Yaowen Zhang, Annanya Sethiya, Adam Takaoglu, Divya Prabhu, Aining Fan, Shravani Vitalapuram, Jurij Dolenšek, Marko Gosak, Richard KP Benninger, Andraž Stožer, and Vira Kravets. Role of GLP1-receptor-mediated α-β-cell communication in functional β-cell heterogeneity. bioRxiv, 2025.

25. Vira Kravets, Claire H. Levitt, and Richard K. Benninger. 195-OR: To Which Degree Do Alpha Cells Shape the Role of the Beta Cells First Responders? Diabetes, 72(Supplement 1):195–OR, June 2023.

26. Michela Riz, Matthias Braun, and Morten Gram Pedersen. Mathematical modeling of heterogeneous electrophysiological responses in human β-cells. PLOS Computational Biology, 10(1):1–14, 01 2014.

27. Daniele Andrean and Morten Gram Pedersen. Machine learning provides insight into models of heterogeneous electrical activity in human beta-cells. Mathematical Biosciences, 354:108927, 2022.

28. Chae Young Cha, Yasuhiko Nakamura, Yukiko Himeno, Jianwu Wang, Shinpei Fujimoto, Nobuya Inagaki, Yung E. Earm, and Akinori Noma. Ionic mechanisms and Ca2+ dynamics underlying the glucose response of pancreatic β cells: a simulation study. J Gen Physiol, 138(1):21–37, July 2011.

29. Vira Kravets, JaeAnn M. Dwulet, Wolfgang E. Schleicher, David J. Hodson, Anna M. Davis, Laura Pyle, Robert A. Piscopio, Maura Sticco-Ivins, and Richard K. P. Benninger. Functional architecture of pancreatic islets identifies a population of first responder cells that drive the first-phase calcium response. PLOS Biology, 20(9):e3001761, September 2022. Publisher: Public Library of Science.

30. Yukari Takeda, Akira Amano, Akinori Noma, Yasuhiko Nakamura, Shimpei Fujimoto, and Nobuya Inagaki. Systems analysis of GLP-1 receptor signaling in pancreatic β-cells. American Journal of Physiology-Cell Physiology, 301(4):C792–C803, October 2011. Publisher: American Physiological Society.

31. Leonid E. Fridlyand and Louis H. Philipson. Pancreatic Beta Cell G-Protein Coupled Receptors and Second Messenger Interactions: A Systems Biology Computational Analysis. PLOS ONE, 11(5):e0152869, May 2016. Publisher: Public Library of Science.

32. Danh-Tai Hoang, Manami Hara, and Junghyo Jo. Design Principles of Pancreatic Islets: Glucose-Dependent Coordination of Hormone Pulses. PLOS ONE, 11(4):e0152446, April 2016. Publisher: Public Library of Science.

33. P. Smolen, J. Rinzel, and A. Sherman. Why pancreatic islets burst but single beta cells do not. the heterogeneity hypothesis. Biophysical Journal, 64(6):1668–1680, 1993.

34. Andraž Stožer, Eva Paradiž Leitgeb, Viljem Pohorec, Jurij Dolenšek, Lidija Križančić Bombek Marko Gosak, and Maša Skelin Klemen. The Role of cAMP in Beta Cell Stimulus–Secretion and Intercellular Coupling. Cells, 10(7):1658, July 2021. Publisher: Multidisciplinary Digital Publishing Institute.

35. Qiang Ni, Ambhighainath Ganesan, Nwe-Nwe Aye-Han, Xinxin Gao, Michael D. Allen, Andre Levchenko, and Jin Zhang. Signaling diversity of PKA achieved via a Ca2+-cAMP-PKA oscillatory circuit. Nat Chem Biol, 7(1):34–40, January 2011. Publisher: Nature Publishing Group.

